# The computational bottleneck of basal ganglia output (and what to do about it)

**DOI:** 10.1101/2024.10.23.619790

**Authors:** Mark D. Humphries

## Abstract

What the basal ganglia do is an oft-asked question; answers range from the selection of actions to the specification of movement to the estimation of time. Here I argue that *how* the basal ganglia do what they do is a less-asked but equally important question. I show that the output regions of the basal ganglia create a stringent computational bottleneck, both structurally, because they have far fewer neurons than do their target regions, and dynamically, because of their tonic, inhibitory output. My proposed solution to this bottleneck is that the activity of an output neuron is setting the weight of a basis function, a function defined by that neuron’s synaptic contacts. I illustrate how this may work in practice, allowing basal ganglia output to shift cortical dynamics and control eye movements via the superior colliculus. This solution can account for troubling issues in our understanding of the basal ganglia: why we see output neurons increasing their activity during behaviour, rather than only decreasing as predicted by theories based on disinhibition, and why the output of the basal ganglia seems to have so many codes squashed into such a tiny region of the brain.

**Significance statement:** The basal ganglia are implicated in an extraordinary range of functions, from action selection to timing, and dysfunctions, from Parkinson’s disease to obsessive compulsive disorder. Yet however the basal ganglia cause these functions and dysfunctions they must do so through a group of neurons that are dwarfed in number by both their inputs and their output targets. Here I lay out this bottleneck problem for basal ganglia computation, and propose a solution to how their outputs can control their many targets. That solution rethinks the output connections of the basal ganglia as a set of basis functions. In doing so, it provides explanations for previously troubling data on basal ganglia output, and strong predictions for how that output controls its targets.

## Introduction

How do the basal ganglia do any useful work? I will argue here that they suffer from a severe computational bottleneck. Their output nuclei, through which they connect with the rest of the brain, are markedly smaller than both their input sources and output targets. Moreover, the standard view is that the basal ganglia’s output nuclei encode by disinhibition, by the cessation of their inhibitory output (Chevalier et al., 1985; Chevalier and Deniau, 1990; Hikosaka et al., 2000; Basso and Sommer, 2011), which provides limited capacity for carrying information. Yet the moment-to-moment dynamics of the basal ganglia are implicated in a lengthy list of proposed functions, including action selection (Redgrave et al., 1999; Klaus et al., 2019), motor program selection (Mink, 1996), kinematic gain control (Turner and Desmurget, 2010; Park et al., 2020), perceptual decision making (Ding and Gold, 2013; Yartsev et al., 2018), duration estimation (Buhusi and Meck, 2005; Gouvêa et al., 2015; Mello et al., 2015; Monteiro et al., 2023), signal routing (Stocco et al., 2010), and more. How one or more such complex functions are enacted through an output signal that has limited capacity in both size and dynamics is unclear. I will begin here by defining this computational bottleneck problem, first detailing the anatomical expansion between the basal ganglia output nuclei and their targets, then arguing that the disinhibition view of basal ganglia output is limited. This sets up two fundamental problems for the basal ganglia output: one, how does it re-expand? And, two, what dynamics does it use to code?

I propose a solution to both these size and coding problems: that the basal ganglia output neuron’s projections to their targets are a set of basis functions, and the output neurons’ activity sets the weights of those functions. All of these ideas will be elaborated below. This solution explains both how basal ganglia output can expand to the same scale as its targets, and why it would need to both decrease and increase its activity. It can also account for troubling features of basal ganglia output, including why it has so many apparently different coding schemes. Consequently, it is a step towards reconciling the basal ganglia’s many apparent functional roles and may shed further light on why dysfunction of the basal ganglia is implicated in so many neural disorders.

### The computational bottleneck problem

#### The structural bottleneck

In rodents, the basal ganglia output nuclei are traditionally considered to be the subtantia nigra pars reticulata (SNr) and entopeduncular nucleus (EP). In primates, the latter is equivalent to the internal segment of the globus pallidus (GPi). Regardless of their names, these all share common anatomical properties: they receive input from the striatum and output to structures including multiple regions of the thalamus, the superior colliculus, and the upper brainstem (Deniau and Chevalier, 1992; McElvain et al., 2021).

The striatum dwarfs the output nuclei. In rats, the striatum in one hemisphere contains around 2.8 million neurons, whereas the SNr and EP combined contain around 30,000 (Oorschot, 1996), smaller by a factor of 100. In mice, the striatum contains about 400,000 projection neurons and the SNr around 12,000 neurons (numbers from the Blue Brain Project Cell Atlas, Rodarie et al., 2022); assuming that half of all projection neurons are D1-expressing and so project to the SNr, this gives a ratio of about 16:1 striatal projection neurons projecting to every SNr neuron in mice. This convergence of striatal projection neurons onto the basal ganglia output nuclei is well known, but we know little about the sizes of the target regions of the output nuclei.

To better understand the scale of the structural bottleneck, I used data from the Allen Mouse Brain Connectivity Atlas (Oh et al., 2014) to first identify a complete set of projection targets of the mouse SNr. The Atlas contains six experiments in which a fluorescent anterograde tracer was injected into the right SNr and filled at least 20% of its volume. For each experiment I found which of a set of 295 non-overlapping target brain regions had evidence of projections from the SNr, by checking if the density of tracer in that region exceeded some threshold. A threshold was necessary to eliminate image noise and other artefacts: without one, all 295 regions contained fluorescent pixels, implausibly implying the SNr projects to every area of grey matter in the mouse brain, from medulla to olfactory bulb (Methods). The number of neurons in each retained target region was found from the Blue Brain Project’s Cell Atlas for the mouse brain (Rodarie et al., 2022). The total number of neurons in SNr target regions scaled with the size of the tracer’s injection volume (Figure 1a). All injection volumes were smaller than the volume of the mouse’s SNr. Fitting a linear model to the scaling let me extrapolate to the number of neurons targeted by the whole SNr (Figure 1a, grey lines), and so estimate the ratio of target neurons to SNr neurons. This ratio fell to a stable value with increasingly stringent thresholds for eliminating noise (Figure 1c), estimated as 154:1.

**Figure 1:**
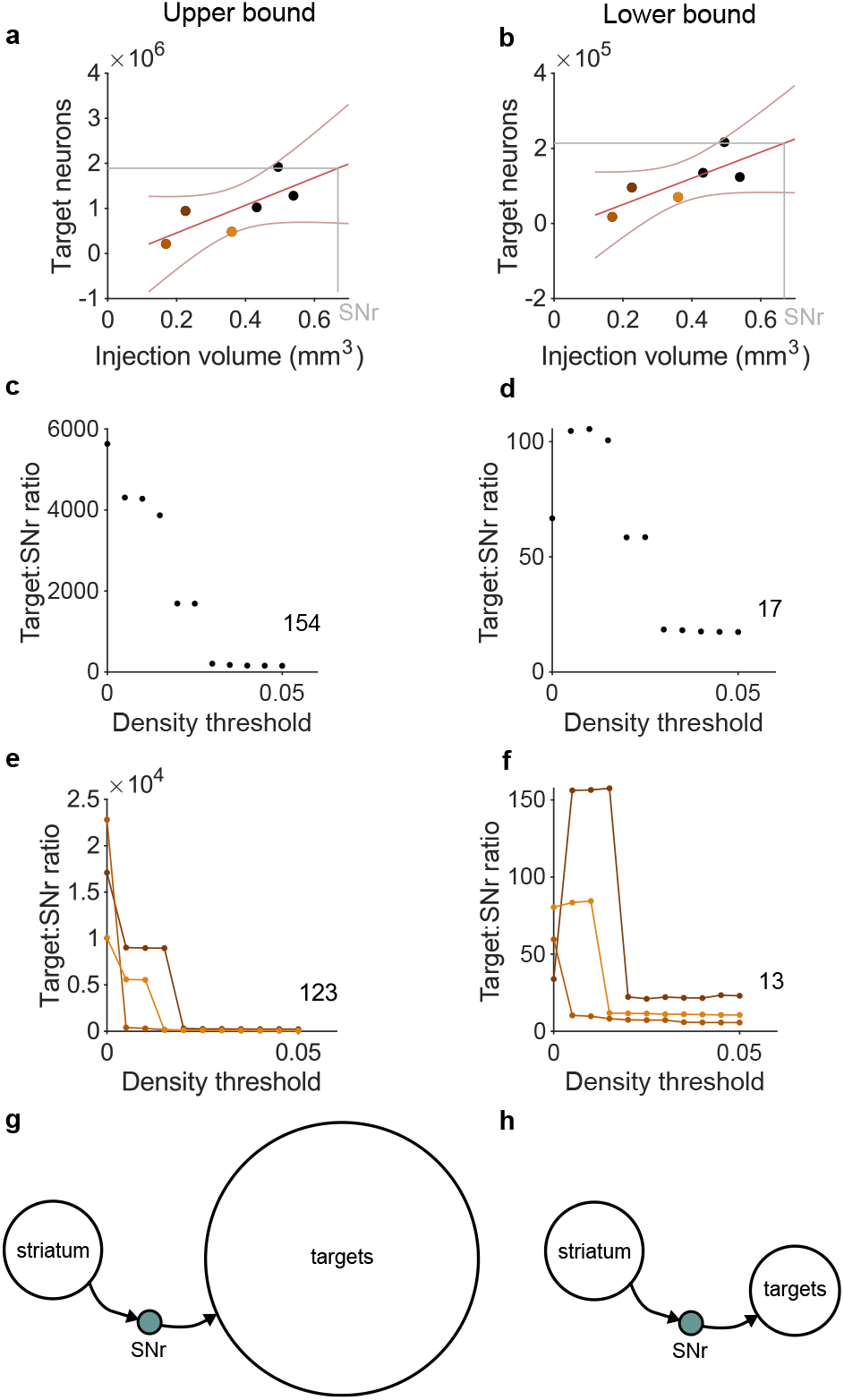
The structural bottleneck of basal ganglia output in the mouse. **a** The total number of neurons in SNr target regions scales with the volume of tracer injection. Each symbol is an estimate from one tracing experiment of the Allen Mouse Brain Atlas; non-black symbols are experiments with more than 90% of the injection within SNr. Red lines show linear fit and 95% confidence interval. Grey line is the extrapolated total neurons in SNr targets from its volume. **b** As panel a, but estimating the targeted neurons in each region from the density of tracer in that region’s volume (Methods). **c** The ratio of target neurons to SNr neurons for a range of thresholds on the minimum tracer density needed to include a target region. Data in panel a is for a threshold of 0.05. Number is the asymptotic estimate of the ratio. **d** As panel c, but estimating the targeted neurons in each region from the density of tracer in that region’s volume (Methods). **e** As panel c, for each of the three tracer experiments with injection volume confined to the SNr (non-black symbols in panel a). Here the expected number of neurons in SNr target regions is computed by scaling that experiment’s total number of target neurons by the proportion of SNr filled by the injection. Number is the asymptotic estimate of the ratio averaged over the three experiments. **f** As panel e, but estimating the targeted neurons in each region from the density of tracer in that region’s volume. **g** Schematic of the upper bound of the SNr bottleneck. Target size from panel e; striatal D1R population estimated at 200,000 neurons (see text). **h** As for panel g, for the lower bound of the SNr bottleneck, target size from panel f.

The extrapolation to the whole SNr’s projection was based on three experiments that had less than 40% of their injection volume inside the SNr (black symbols in Figure 1a), potentially including neighbouring regions of the SNr that have different connection patterns. Weighting the linear model fit by the proportion of the injection inside the SNr gave practically identical ratios of target to SNr neurons (not shown). Estimating the number of target neurons directly for each of the three tracer injections almost wholly within the SNr (non-black symbols in Figure 1a), by extrapolating from the volume of their injection (Methods), also resulted in similar and stable ratios for the total number of target neurons to the number of SNr neurons (Figure 1e).

The total number of neurons in the target regions is an upper bound on the number of connections made by SNr neurons. To estimate a lower bound, I approximated the arborisation of the axons from SNr in the target region by the volume density of the tracer in that region, scaling the number of neurons in each target region by the proportion of its volume occupied by the tracer (Methods). This lower bound estimate of target neurons also scaled with the tracer’s injection volume (Figure 1b), reached a stable value with increasing noise threshold (Figure 1e), and was robust to alternative calculation using the three within-SNr experiments (Figure 1f).

The estimated expansion from the basal ganglia output nucleus SNr to its targets ranges from about 1:154 down to 1:13 (Figure 1). Even the lower bound on this expansion is thus about as large as the compression of inputs from striatum: the basal ganglia’s output is then a considerable bottleneck, compressing its inputs by at least 10:1, and re-expanding them in its output targets by at least 1:10 (Figure 1g-h).

#### The dynamic bottleneck (or, why not disinhibition)

Basal ganglia output neurons are constantly active. In rodents, they typically average 30 spikes/s; in primates, around 60 spikes/s. They are also all GABAergic. This constant stream of high-frequency GABA release on to their target neurons has naturally led to the assumption that they constantly inhibit their targets. From that has followed the disinhibition hypothesis (Chevalier et al., 1985; Deniau and Chevalier, 1985; Chevalier and Deniau, 1990) that releasing this inhibition is key to how the basal ganglia encode information, by allowing their target neurons to respond to their inputs. This signalling by disinhibition is the basis for most prominent conceptual (Mink, 1996; Redgrave et al., 1999; Hikosaka et al., 2000) and computational (Gurney et al., 2001; Frank, 2005; Humphries et al., 2006; Leblois et al., 2006; Bogacz and Gurney, 2007; Vitay and Hamker, 2010; Liénard and Girard, 2014; Lindahl and Hellgren Kotaleski, 2016; Dunovan et al., 2019) models of the basal ganglia.

Yet it is a myopic view of basal ganglia output that places a strong limitation on the potential dynamic range of the output neurons, allowing coding by only a decrease in activity, and then often reduced to just whether the activity is on or off, a binary signal (e.g. Hikosaka et al., 2000). There is no role for increases in activity, or changes to the patterns of activity (there are other theories: for example, there is evidence that the output of the basal ganglia in songbirds controls the timing of activity in their thalamic targets Goldberg et al. 2013). This compounds the structural bottleneck by then further limiting how each neuron can send information. For example, taking the binary on/off signal literally, disinhibition reduces the information coding capacity of the whole basal ganglia output to just one bit per neuron, a few thousand bits in total.

### A solution: basal ganglia outputs are dynamic weights

The basal ganglia’s computational bottleneck problem is, then, that we have a limited number of free parameters – the output neurons and their dynamical range – compared to the number of outputs that we need to control. My proposed solution to this problem is a reframing of what the basal ganglia output encodes.

I propose that the basal ganglia’s output connections are best understood as basis functions, and the level of basal ganglia output activity sets the weights on those functions. Let’s unpack those ideas, starting with a definition of basis functions.

Basis functions are a typical solution for how to use a few parameters to control a large range of output. Figure 2a shows the key ideas. We first tile the output range we want to control with a set of basis functions, such as the five Gaussians in the example of Figure 2a. Each basis function has a single weight that sets its contribution, such as the amplitude of a Gaussian (Figure 2a, middle). Summing basis functions of different weights can then create many different output functions over a large range of outputs (Figure 2a, bottom). Basis functions thus create an expansion from a few controllable parameters – the weights – to a much larger target space.

**Figure 2:**
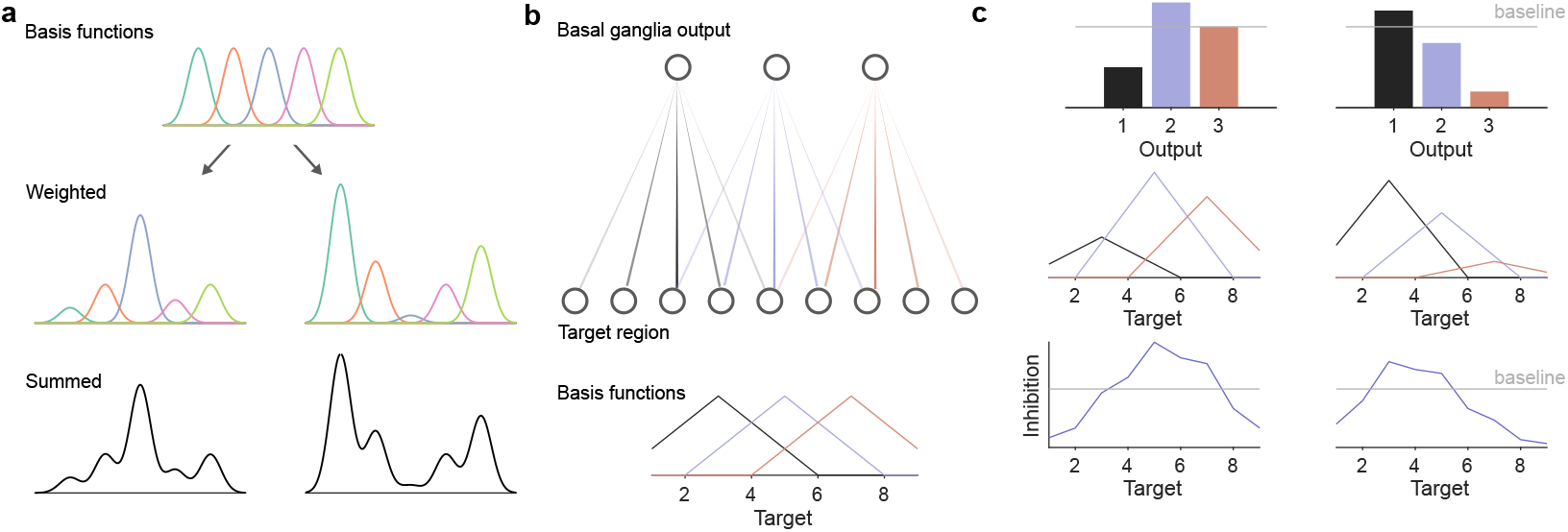
Basis functions and basal ganglia output. **a** Schematic of basis functions. A range of values is tiled by a set of basis functions, here five Gaussians (top). Each basis function’s contribution is controlled by a single weight: the middle panels show two different weightings. Summing these weighted basis functions creates a continuous function spanning the range of values (bottom), controlled by just five parameters, the weights on the basis functions. **b** Basal ganglia output connections define basis functions. Top: idealised network showing distribution of basis function strengths, fanning from the output nuclei to a larger target region; colour intensity is proportional to strength. Bottom: the basis functions created by the connection strengths. **c** Basal ganglia output activity parameterises the basis functions. Using the network in panel b, two examples of how basal ganglia output activity (top) scales each neuron’s basis function (middle), which when summed as the input to each target creates a continuous inhibition function (bottom). Grey line indicates baseline output activity (top) and consequent inhibition of targets (bottom).

Now consider that the connections of the basal ganglia output neurons will have a distribution of strengths (Figure 2b, top), the strength of a connection between an output neuron and its target being the product of the number of synapses and the conductances of those synapses. The idea then is that this distribution of strengths defines the basis functions (Figure 2b, bottom).

Consequently, the amplitude of basal ganglia output activity sets the weights of those basis functions. Figure 2c shows two examples of what this would look like: a particular vector of basal ganglia output activity scales the basis functions created by the output strengths; when summed at each target, the larger target region as a whole receives a continuous function of inhibition, specified by far fewer basal ganglia output neurons than target neurons.

This idea rests on just two assumptions: that connections of the output neurons have a distribution of strengths, and that these distributions overlap. It seems vanishingly unlikely that a given output neuron has an identical effective influence on each of its target neurons, so a distribution of strengths seems reasonable. And because the output of the basal ganglia is topographically organised (Deniau and Chevalier, 1992; Hoover and Strick, 1993; Lee et al., 2020; Foster et al., 2021; McElvain et al., 2021), with adjacent neurons projecting to adjacent targets, we might also reasonably expect these distributions of strengths to overlap. Beyond that, I emphasise that the schematics in Figure 2 are for illustration, not theory: I’m not claiming that distributions of output strengths have to be symmetric, nor that their “centres” are distributed equidistant from each other in some topographic space, nor that the distributions have to be the same, nor that they have to follow any specific basis function used in the literature (such as radial basis functions). Rather, the theory proposed is perhaps best expressed as: basal ganglia output activity is a dynamic weight on some function defined by the strengths of the output connections.

Let’s now state the most general form of the theory and derive some general predictions from it.

### The general form of the theory and its predictions

Consider that *b* basal ganglia output neurons project to a set of *n* target neurons. We have already established that *b < n*, the structural bottleneck. The theory proposes that the goal of basal ganglia output is to create a specific function of inhibition across those *n* target neurons, which we can describe in a *n*-dimensional vector **f** with entries *fi* ≤ 0.

The theory can thus be expressed as the linear system

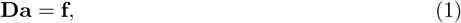

where **a** is the *b*-dimensional vector of basal ganglia output activity, and **D** is the *n* × *b* matrix of connection weights from the basal ganglia output to the set of target neurons (Figure 3a). Their values are also constrained: *ai* ≥ 0 as neural activity cannot be negative, and *Dij* ≤ 0 because basal ganglia output is inhibitory. Matrix **D** defines the basis functions, one column per output neuron. For example, in the schematic model of Figure 2b each column of **D** is a shifted version of the same, symmetric basis function. Thus the vector **a** of basal ganglia activity are the dynamic weights on the basis functions **D** that gives the target function **f**.

**Figure 3:**
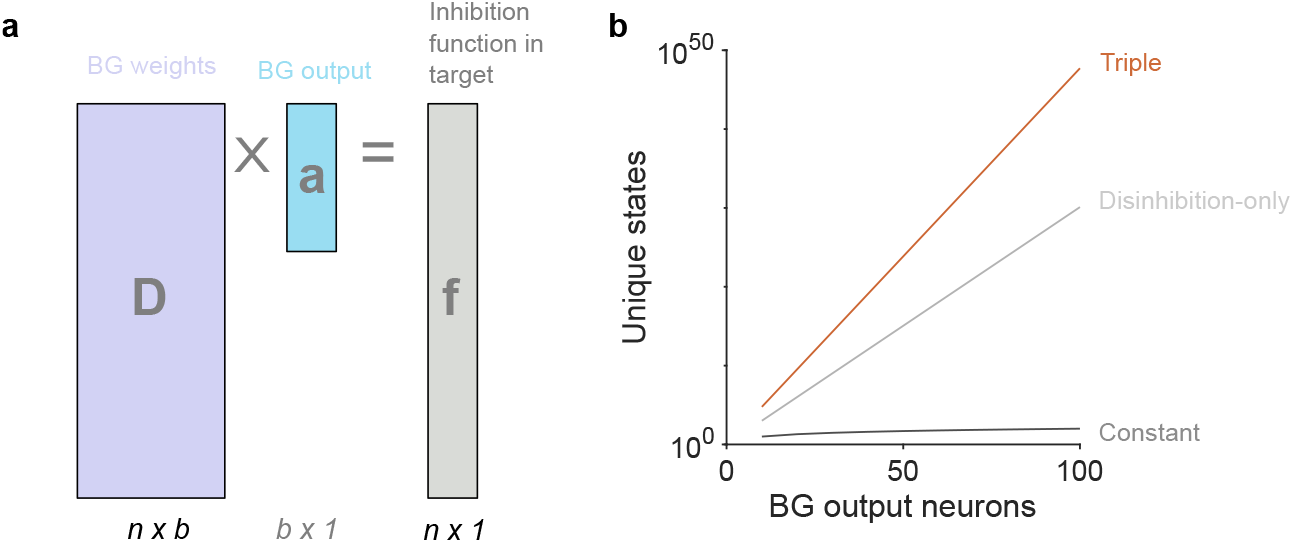
Linear system model of basal ganglia output. **a** Schematic of the linear model in Eq. 1. **b** Scaling of the number of possible functions **f** defined by basal ganglia outputs. Each line give the number of unique functions possible with that number *b* of output neurons. Different lines correspond to the different number of states each output neuron can meaningfully be in: constant (1); disinhibition (2: on/off); triple (3^*b*^) is up/down/unchanged.

#### Prediction of non-uniform inhibition from uniform output

Basal ganglia output neurons **a** are tonically active, firing at rest at a rate of around 30 spikes/s in rodents and 60 spikes/s in primates. That **a** has non-zero values means a non-trivial target function **f** is always defined.

However, many theories implicitly assume that this tonic firing necessarily means there is a uniform level of inhibition, such that all values of **f** are the same. This is implied by theories that tonic inhibition defines the “no-go” or “off” signal for selecting responses, actions, or motor programs (Mink, 1996; Redgrave et al., 1999; Hikosaka et al., 2000). The model in Eq. 1 shows this is only true if the rows of **D** have the same sum. But this is unlikely as only the columns of **D**, being the projections of each output neuron, are defined by development and plasticity. Consequently, the model predicts that tonic activity of a set of output neurons causes a non-uniform inhibition of their targets.

#### Prediction of increased output activity

Traditionally, it is the cessation of this tonic inhibition, the disinhibition, that has been the basis for key conceptual (Mink, 1996; Redgrave et al., 1999; Hikosaka et al., 2000) and computational (Gurney et al., 2001; Frank, 2005; Humphries et al., 2006; Leblois et al., 2006; Bogacz and Gurney, 2007; Vitay and Hamker, 2010; Liénard and Girard, 2014; Lindahl and Hellgren Kotaleski, 2016; Dunovan et al., 2019) models of the basal ganglia’s function.

The model offered in Eq. 1 places no constraints on the values of **a** around the baseline tonic activity. Rather, the tonic activity values of **a** define a default **f** from which behaviourally-necessary changes to **f** must occur. This predicts that both increases as well as decreases in output neuron activity can change **f** when basal ganglia output must cause or influence some behavioural event.

This prediction is borne out by data. Basal ganglia output neuron activity does increase in many tasks (Gulley et al., 1999; Handel and Glimcher, 1999; Gulley et al., 2002; Sato and Hikosaka, 2002; Jin and Costa, 2010; Fan et al., 2012; Rossi et al., 2016; Schwab et al., 2020) and the increases are as equally time-locked to action as the decreases (Sato and Hikosaka, 2002; Jin and Costa, 2010; Fan et al., 2012). In some reports output neuron activity can seemingly continuously encode parameters both above and below the nominal “tonic” firing rate (Barter et al., 2015).

Allowing for increased output activity increases the basal ganglia’s scope for control. Restricting ourselves to classic disinhibition allows just two output states, on and off. The number of possible unique output combinations is then 2^*b*^ (Figure 3b). But adding just one more output state, the increase above the tonic level, makes the number of unique input combinations 3^*b*^ (Figure 3b): at 100 output neurons, this triple state can achieve more then 5 × 10^47^ unique dynamic weight combinations, and hence that many different functions of inhibition **f**. Consequently, even a small group of basal ganglia output neurons could control a wide repertoire of states in its target structures.

#### Prediction of low variability in output activity

Stating the theory as the linear system Eq. 1 lets us ask an interesting question: is basal ganglia output degenerate? That is, in some behavioural event for which the basal ganglia are necessary, can different combinations of increases and decreases of output neuron activity achieve the same behavioural effect?

Let’s assume that the same behavioural effect means achieving the same target function **f**. Then we are asking how many solutions exist to Eq. 1 (Druckmann and Chklovskii, 2012): how many different basal ganglia outputs **a** achieve the same target function **f**.

A heterogeneous linear system like Eq. 1 can have no, one, or an infinite number of such solutions. As we are interested in events where the basal ganglia have a necessary role, then by definition we are interested in the set of **f** that can be achieved by the basal ganglia output given **D**. So there must be at least one solution **a** for a given, behaviourally-relevant, target function **f**. But is there more than one?

It seems unlikely. This is because the matrix of connection weights **D** is almost certainly full rank, having no linearly dependent columns. For structured basis functions, where each column of **D** is approximately a shifted version of the same function (like Figure 2a), we can guarantee that **D** is full rank by construction (Methods). For random basis functions, selecting the values of **D** from a wide range of symmetric probability distributions would guarantee it was full rank (Rudelson and Vershynin, 2009). A linear system with full rank **D** has at most one solution. Thus, either the basal ganglia output connections have a genetically defined low-rank structure or there is only one basal ganglia output **a** that can achieve a given target function of inhibition **f**.

Having exactly one solution **a** to Eq. (1) predicts that individual basal ganglia neurons would show little variability between repeated behavioural events that need the same **f**. I am proposing here that the necessary solution to **a** is for each behavioural event, so the predicted time-scale of this variation is around the gross changes in firing rate time-locked to an event, not precise spike-timing.

Conversely, observing considerable variability in basal ganglia output activity between repeated events would imply either that **f** is not the goal of basal ganglia output, so the theory here is incorrect, or that the connections of the basal ganglia to their targets are linearly dependent and so basal ganglia output is redundant. This in turn would imply strong constraints on how basal ganglia output is wired, in order to achieve this linear dependence.

I am unaware of convincing data either way. While there is a considerable literature on single neuron activity in both rodent SNr (e.g. Gulley et al., 1999, 2002; Bryden et al., 2011; Fan et al., 2012) and primate GPi/SNr (e.g. Handel and Glimcher, 1999; Sato and Hikosaka, 2002; Nevet et al., 2007; Sheth et al., 2011) during tasks, I am not aware of any that have quantified the trial-to-trial variability in that activity during exact repetition of a behaviour for which the basal ganglia are necessary. Close examination of example raster plots of individual SNr neurons aligned to the onset of eye movements (e.g. Handel and Glimcher, 1999; Sato and Hikosaka, 2002) suggests that gross changes of activity are highly consistent between trials, unlike, say, the rate variation of individual cortical neurons between repeated sensory stimuli (Tolhurst et al., 1983).

### How the solution could work in practice

Let’s illustrate the idea of basal ganglia outputs as dynamic weights in two concrete instantiations: the control of cortical state by basal ganglia output to thalamus; and the control of superior colliculus’ coding of saccade target by its inputs from the SNr.

#### Basal ganglia output control of the repertoire of cortical states

The basal ganglia’s output to the thalamus is the main focus of much theorising because of its potential to control the dynamics of the cortical targets of those thalamic regions (e.g. Humphries and Gurney, 2002; Frank, 2005; Dunovan et al., 2019; Möller and Bogacz, 2019; Athalye et al., 2020; Logiaco et al., 2021). But the thalamic regions contain more neurons than the basal ganglia output nuclei, and the thalamus in turn is dwarfed by the numbers of cortical neurons (consider, for example, that of all the synapses arriving onto layer 4 neurons in the visual cortex, thalamic synapses make up just 5% of the total Peters and Payne 1993). Here I illustrate how the idea of dynamic weights defined by the combinations of a few basal ganglia output neurons can allow control of cortical dynamics. In general, the problem of how a few inputs can drives the states of a larger dynamical system is studied under the control theory concept of *controllability* (Sontag and Sussmann, 1997; Liu et al., 2011; Gu et al., 2015; Kao and Hennequin, 2019); an interesting extension of the work here would use controllability approaches to identify what form of target function **f** is ultimately necessary to control cortical states, and which elements of the cortical circuit must be targeted to do so.

Let’s consider a recurrent neural network (RNN) to model a region of cortex, as these nicely capture the basic problem: a network of mixed excitatory and inhibitory neurons that is capable of producing complex dynamics (Figure 4a). A brief step in input to this network produces a population response (Figure 4b, top). We can characterise this population response by the trajectory it creates in a low-dimensional space (Figure 4b, bottom). Such trajectories of neural activity in cortex correspond to specific arm movements (Churchland et al., 2012; Gallego et al., 2017; Rodriguez et al., 2024), elapsed durations (Voitov and Mrsic-Flogel, 2022), or choices (Harvey et al., 2012).

**Figure 4:**
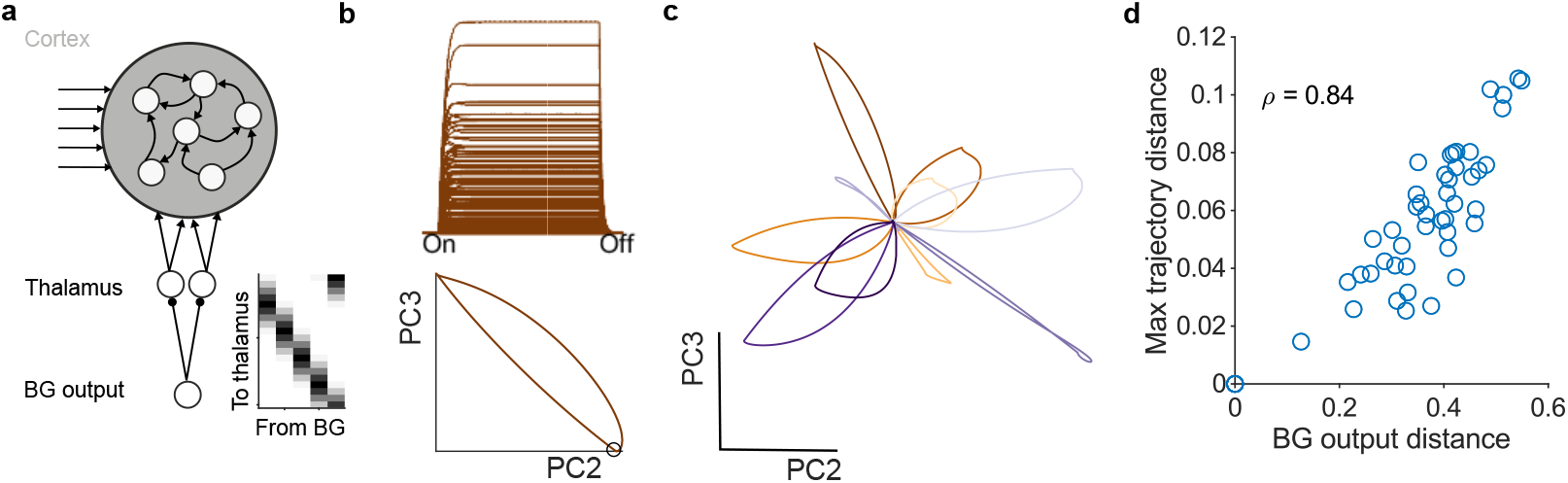
Basal ganglia output control of cortical dynamics. **a** Schematic of the recurrent neural network (RNN) model, its inputs from other cortical regions (arrows) and its inputs from thalamus. Basal ganglia output to thalamus is a set of overlapping symmetric basis functions (inset; grey-scale indicates strength of connection, white indicates no connection). Thalamus projects to 10% of the cortical RNN units. Example simulations use 5 basal ganglia outputs, 20 thalamic units, and 200 cortical units. **b** Example response of all RNN units to stepped input (top), and projection of that RNN activity into a low-dimensional space (PC: principal component). The trajectory of low-dimensional activity captures the move away from and return to baseline activity (black dot). **c** Trajectories of RNN activity in response to different basal ganglia outputs. Each line plots the trajectory in response to a different basal ganglia output vector, with all other inputs held constant. Output vectors were sampled from a uniform distribution centered on tonic activity, modelling both increases and decreases of output. **d** Variation in basal ganglia output maps to variation in the trajectories of RNN activity. *ρ*: Spearman’s rank.

A small fraction of these inputs, 10%, are from the thalamus. These thalamic inputs are in turn controlled by a small handful of basal ganglia output neurons, whose outputs create a set of basis functions to control thalamic activity. These basis functions are symmetric, overlapping, and tiled in a ring as shown in Figure 4a (inset). The Appendix discusses the constraints on the number of thalamic neurons and basal ganglia weight distributions implied by this model.

Despite the small number of basal ganglia outputs they are sufficient to qualitatively change the state of the cortical circuit. With all other inputs held the same, different vectors of basal ganglia output create different trajectories of activity in cortex (Figure 4c). They do so by creating different functions of inhibition in the thalamus, defined by both decreases (disinhibition) and increases in basal ganglia output activity. Crucially, we see that these changes in trajectory would alter cortical coding (of an arm movement, a delay, a choice) without need for that coding in the basal ganglia output.

Work in songbirds has argued that basal ganglia activity is necessary to explore the repertoire of potential movements, including the variation of syllable generation in songs (Ölveczky et al., 2005; Singh Alvarado et al., 2021). The dynamic weights view is also consistent with this argument: the trajectories of activity in the cortical circuit vary in proportion to the distance between the output vectors of the basal ganglia (Figure 4d). So greater variation in basal ganglia output could map to greater exploration of the cortical repertoire of dynamics, and thus to motor behaviour or choices (de A Marcelino et al., 2023).

#### How output neurons can control superior collicular activity to influence the orientation of the eyes and body

Another major target of basal ganglia output is the intermediate layers of the superior colliculus. This structure plays a key role in orienting the eyes and body (Hikosaka et al., 2000; Felsen and Mainen, 2008; Villalobos and Basso, 2022), with activity in its intermediate layer acting as a command signal for eye movements to a particular location (Hikosaka et al., 2000). Much ink has thus been spilt on how the inhibitory signals emanating from the basal ganglia to the superior colliculus may in turn control the direction of gaze (Hikosaka and Wurtz, 1985; Jiang et al., 2003; Girard and Berthoz, 2005; Basso and Sommer, 2011).

Most theories agree on the following (Hikosaka et al., 2000; Jiang et al., 2003; Girard and Berthoz, 2005; Chambers et al., 2011; Thurat et al., 2015). The intermediate, or motor, layer of the superior colliculus represent the direction of gaze in two-dimensional retinotopic co-ordinates. For convenience we’ll consider them as a Cartesian grid of (x,y) positions in a two-dimensional plane. Neural activity in the intermediate layer thus represents a motor command to direct gaze towards the location represented by the active neurons (Lee et al., 1988; Anderson et al., 1998). Basal ganglia inhibition of these collicular neurons is able to suppress changes in gaze direction to remembered locations (Mahamed et al., 2014) and possibly stimulus-driven locations. Consequently, a pause in basal ganglia output directed at collicular neurons representing a particular location allows the change in gaze to happen (Hikosaka and Wurtz, 1985).

Less clear is how the basal ganglia output can provide that fine control over neural activity in the colliculus. The straightforward solution (Dominey and Arbib, 1992; Dominey et al., 1995; Jiang et al., 2003; Girard and Berthoz, 2005) is that the basal ganglia output also has a two-dimensional retinotopic map, and hence provides point-to-point control over collicular activity (Fig. 5a, top). But this scheme scales poorly because the number of possible co-ordinates scales linearly with the number of neurons (Fig. 5b, blue). And it seems at odds with the few neurons involved: of the basal ganglia output nuclei, only the substantia nigra pars reticulata (SNr) projects to the superior colliculus; that projection originates from at most two-thirds of the SNr (Deniau and Chevalier, 1992; McElvain et al., 2021); and even within that region the SNr neurons projecting to the superior colliculus are potentially in the minority, as antidromic stimulation of the colliculus activates far less than half of all sampled SNr neurons (Hikosaka and Wurtz, 1983c; Jiang et al., 2003).

**Figure 5:**
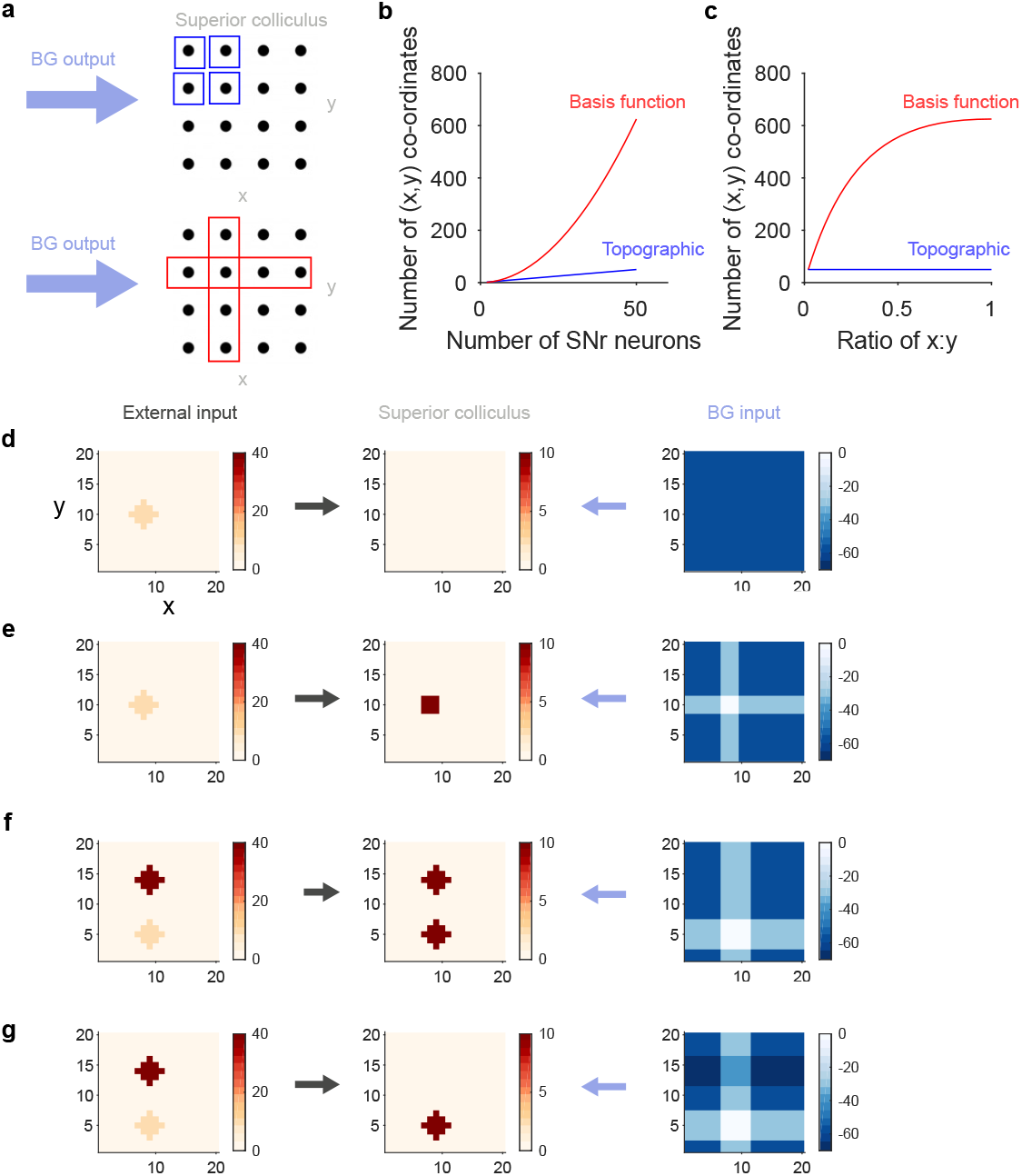
Basal ganglia output control of superior colliculus. **a** Potential schemes for basal ganglia inhibition of saccade targets in the intermediate layers of superior colliculus. The black circles are collicular neurons representing saccade targets in Cartesian co-ordinates. Top: topographic mapping of basal ganglia output to superior colliuclus, one output neuron per co-ordinate (blue). Bottom: a spanning code created by basis functions, with one output neuron’s projection spanning one row or column (red). **b** Scaling of the number of controllable Cartesian co-ordinates by topographic or basis function mapping of basal ganglia output to superior colliculus. Topographic mapping is the best-case scenario of 1 neuron per co-ordinate. **c** Scaling of controllable co-ordinates with grid asymmetry. For a fixed number (50) of basal ganglia output neurons, the scaling of the number of controllable co-ordinates as the superior colliculus grid moves from a single row to a symmetric grid. **d** Simulations of basis function control of saccadic activity in superior colliculus. External input via the superficial layers specifies a saccade target in Cartesian co-ordinates (left), input to a 20 × 20 grid of collicular neurons middle. Twenty basal ganglia output neurons per side provide tonic inhibition of the superior colliculus: we plot here the inhibition received by each superior colliculus neuron (right). Unchanging inhibition prevents the build-up of saccadic activity at the target location (middle). **e** As for panel d, but now the basal ganglia neurons whose basis functions include the x-coordinates or the y-coordinates of the target location pause their firing, thus allowing at their intersection (right) the build up of saccadic activity (middle). **f** Two competing external inputs (left) could cause saccadic activity to increase at both locations in colliculus (middle), even if only a few basal ganglia output neurons paused their firing (right), because the second, upper target falls in the column covered by paused basal ganglia neurons whose basis functions are the y-coordinates. **g** As for panel f, but with other basal ganglia neurons increasing their firing and hence inhibition of collicular neurons (right, darkest rows), thus suppressing the build-up of saccadic activity (middle) at the second target location.

The basis function idea provides a different solution, of the output neurons defining weights on basis functions tiling the two-dimensional plane. I give one example of how this solution might work; others are possible. In this *spanning code*, the projection of each basal ganglia output neuron (or group of) is a basis function that spans one row or one column of the two-dimensional co-ordinates for gaze direction. Then a particular co-ordinate is specified by the overlapping output of just two neurons (Fig. 5a, bottom).

The spanning code scales well, with the number of controllable co-ordinates rising quadratically with the number of neurons (Fig. 5b, red). As the spanning code uses just two output neurons to signal a particular location, its scaling is slower than a purely combinatorial code (Fig. 3b), and its scaling is slower when the grid is asymmetric (Fig. 5c). But it still scales better than standard theories of basal ganglia output to superior colliculus: for a given number of output neurons there are always more controllable coordinates for the spanning code than for the point-to-point wiring of a topographic map (Fig. 5b-c). So let’s check that the spanning code can indeed control collicular activity to provide appropriate motor commands for gaze direction.

Imagine a model where input specifying the target gaze direction (from e.g. the frontal eye fields or the superficial layer of the superior colliculus) arrives at a grid of intermediate layer collicular neurons (Fig 5d; Methods). Activity at a location on that grid would represent a motor command to shift gaze to that target. At the same time, these neurons receive constant inhibitory input from a set of basal ganglia neurons, each of whose output spans rows or columns of the collicular grid as in Figure 5a. This constant inhibition suppresses all response to the target input (Fig 5d), preventing a shift in gaze direction. Dropping the activity of basal ganglia output neurons whose projections intersect at the target location results in a hill of activity in the intermediate layer in neurons representing that location (Fig 5e). Basis functions can thus allow suppression and selection of gaze direction changes.

This selection requires only a decrease in basal ganglia output, but I have been arguing that they encode bidirectional “dynamic weights”: what then might an increase in basal ganglia output encode here? One answer is to correct for unwanted loss of inhibition elsewhere on the two-dimensional map of gaze directions. Consider two competing target locations that lie on the same column (Fig 5f, left): pausing intersecting basal ganglia outputs for one target could now result in a hill of activity at both target locations on the superior colliculus’ map (Fig 5f, implying two simultaneous but different changes in gaze direction. However, increasing the output of basal ganglia neurons whose basis functions are the corresponding row of the unwanted target location will suppress the hill of activity at this location (Fig 5g). Increasing basal ganglia output could ensure that at most one target location for gaze direction becomes active in the intermediate superior colliculus.

## Discussion

Basal ganglia output neurons are vastly outnumbered by their target neurons in the thalamus and brainstem. To this structural bottleneck is added the further dynamical bottleneck that this output is both constant and inhibitory. It is unclear how the basal ganglia’s output re-expands, both structurally and dynamically, to provide suitable control over its target regions.

I’ve offered as a solution the following idea: the activity of a basal ganglia output neuron defines the weight on a basis function defined by the pattern of its connections to its target regions. In this way, basal ganglia output can re-expand, for arbitrarily complex functions can in principle be constructed from the summation of a set of basis functions. The complex function approximated in this model is the level of inhibition across the target neurons of the basal ganglia output.

This model makes three predictions. First, output neurons all in their tonic state of constant firing do not necessarily imply uniform inhibition of their targets. Second, that both decreases and increases of output neuron firing are necessary to create the target function. Third, that individual output neurons will show low variation in their activity between identical events.

### Implications for our understanding of the basal ganglia

There are three main implications of these ideas and their predictions.

The first is that disinhibition is not the key to how basal ganglia output operates. Weights need to change in both directions to control basis functions, and create the desired target function by their summation. So under this account we’d expect basal ganglia output to both increase or decrease as necessary to set the desired target function of inhibition in its target. This provides a functional explanation for why basal ganglia output activity increases as well as decreases during movement or action (Gulley et al., 1999, 2002; Jin and Costa, 2010; Fan et al., 2012; Barter et al., 2015; Rossi et al., 2016; Schwab et al., 2020), which cannot be accounted for by classic disinhibition-based theories. The second is that we can think of basal ganglia output as allowing rapid exploration of different activity patterns in a target region, which could be crucial in learning; for example of new actions such as skilled limb movements or songs. This is why I’ve called the basal ganglia output “dynamic weights” throughout, as simply by changing their activity the output neurons define a new function of inhibition in their target, which creates a different response in that target to the same input (Fig 4).

The third is that basal ganglia outputs have no intrinsic “code”, but are control signals for a target region. The SNr has many apparent codes. Changes in SNr activity align to the onset of changes in gaze direction in monkeys that are stimulus or memory-driven (Hikosaka and Wurtz, 1983a,b; Handel and Glimcher, 1999, 2000; Basso and Wurtz, 2002; Sato and Hikosaka, 2002) and changes in movement in rodents (Lintz and Felsen, 2016). Jin and Costa (2010) report that SNr activity changes at the beginning and end of an action sequence, not at each action (though such discrete stop and start coding has not been found in striatum Sales-Carbonell et al. 2018). By contrast, Rossi et al. (2016) report SNr activity changes align to each individual lick a mouse makes on a spout. Fan et al. (2012) demonstrate that a pause in SNr activity can last for as long as a mouse holds down a lever is held, seemingly coding duration or a sustained action, rather than action onset or offset. And Barter et al. (2015) offer a startling demonstration of apparently continuous coding of a mouse’s head position in the (x,y) plane by the activity of individual SNr neurons, which typically coded either the x- or y-axis displacement of the head from its central position. Worse, as they are the bottleneck between striatum and the rest of the brain the basal ganglia output nuclei likely inherit other variables known to be encoded in striatum, and there are many of those, including time (Gouvêa et al., 2015; Mello et al., 2015), decision variables (Ding and Gold, 2013; Yartsev et al., 2018), possible actions (Klaus et al., 2019), their predicted value (Samejima et al. 2005; but see Elber-Dorozko and Loewenstein 2018), and their kinematics (Rueda-Orozco and Robbe, 2015; Yttri and Dudman, 2016). How so few neurons could seemingly encode such a range of different variables is unclear. A starting point for a solution could be the dynamic weights theory offered here: basal ganglia output is not coding these variables, but are control signals for encoded variables in their target regions.

### Predictions for effects in target brain regions

The interpretation of basal ganglia output as encoding “dynamic weights” is a general principle for how that output can do useful work in its many target regions of the brain. Further specific predictions of this idea depend on the region targeted.

One prediction is that this re-expansion allows the basal ganglia output to shift cortical activity across an extensive repertoire of different states, via the shaping of thalamic output to cortex. The idea that basal ganglia output shifts the state or trajectory of cortical activity has been gaining traction: Athalye et al. (2020) looked at the problem of how motor cortical activity re-enters a desired state after a reward, and proposed a conceptual model for how plasticity at cortico-striatal synapses may alter basal ganglia output to shift cortical activity; Logiaco et al. (2021) looked at the problem of how motor cortex can drive sequences of movements, developing and analysing a model of how shifting basal ganglia output can switch motor cortical activity states and thus create movement sequences (though based only on disinhibition setting thalamic units on or off). Which regions of cortex this prediction extends too is unclear; speculatively, it would be in the layers (II/III and V) of cortical regions that receive direct input from the motor and intralaminar thalamic nuclei that the basal ganglia output directly influences (Nishimura et al., 1997; Kha et al., 2001; Middleton and Strick, 2002; Bodor et al., 2008; Kuramoto et al., 2011). This prediction is consistent with data from primates (Sauerbrei et al., 2020), rodents (Inagaki et al., 2022) and songbirds (Moll et al., 2023) showing that motor thalamic input to motor cortex is necessary for initiating, or changing to, a discrete element of movement. It follows that manipulating the state of basal ganglia output projections to thalamus should change the state of cortical activity, and hence alter behaviour. Recent studies that optogenetically manipulated SNr axon terminals in motor thalamus reported that inhibiting these SNr terminals during a licking task biased licking to the contralateral side, whereas activating these terminals stopped licking all together (Morrissette et al., 2019) or prevented impulsive licking (Catanese and Jaeger, 2021). These data are consistent with shifting the state of cortical activity; but they are also consistent with classic disinhibition ideas that pauses in basal ganglia output are a go signal and increases in the output are a stop signal. Such optogenetic stimulation that broadly targets all terminals with a stereotyped pattern of stimulation is repeatedly setting all the dynamic weights to approximately the same values. Distinguishing classic disinhibition and dynamic weight accounts of basal ganglia output would instead need selective stimulation of non-overlapping sets of SNr terminals: the dynamic weights idea predicts this would result in different states of cortical activity, and hence potentially different behavioural responses.

If the basis functions defined by the connections of basal ganglia output neurons have some topographic organisation, as in the example of superior colliculus, then different predictions arise for manipulating their activity. Selective optogenetic activation of a few SNr neurons that project to the superior colliculus would be predicted to maximise the weight on their basis functions, strongly inhibiting a region of the intermediate superior colliculus, and so create a region of visual space that it is difficult to shift gaze or orient towards. The shape of that region would depend on the shape of the basis functions. The specific spanning code advanced here (Fig 5) would predict that the region of visual space would cover entire x- or y- co-ordinates. This prediction is consistent with reports that the activity of individual SNr neurons encodes the entire x-axis or y-axis displacement of the head in mice (Barter et al., 2015). That said, as noted above the spanning code is not an optimal form of basis function tiling, and other tilings of the collicular representation of gaze direction, including random projections, would be worth further exploring.

### What the dynamic weights idea does not yet address

There are a few key issues that further development of this “dynamic weights” idea could usefully address. The first is its precise role in behaviour. Most current basal ganglia theories focus on how its output controls action, but fall largely into two opposing camps.

One is that basal ganglia output controls action selection, via pauses in its tonic inhibition (Redgrave et al., 1999; Bogacz and Gurney, 2007; Liénard and Girard, 2014; Klaus et al., 2019); the other is that basal ganglia output controls action specification, by its reduced activity modulating some aspect of the kinematics or gain of movement (Turner and Desmurget, 2010; Park et al., 2020; Thura et al., 2022). The dynamic weights idea is currently agnostic to these theories: indeed, it suggests that coding is in the target region, not in basal ganglia output per se. Changes in cortical state by basal ganglia output could be the selection of action or change in the kinematics of an action. Similarly, the example given of superior colliculus control follows previous models of SNr output to colliculus (Girard and Berthoz, 2005) in interpreting the hill of activity as selecting an action, the new direction of gaze; but one could interpret this as specifying action, setting the velocity of the change in gaze direction by controlling the size of the hill of activity.

Both canonical action selection (e.g. Gurney et al., 2001; Frank, 2005; Humphries et al., 2006; Liénard and Girard, 2014; Lindahl and Hellgren Kotaleski, 2016) and specification (e.g. Yttri and Dudman, 2016; Park et al., 2020) theories also assume basal ganglia output are organised as a collection of parallel populations or “channels”, each one representing an option to be selected or specified. While this assumption is the basis for powerful explanations of how the basal ganglia circuit can implement selection or specification, it also strongly restricts the coding capacity of the basal ganglia output. The theory here is as yet agnostic to the size of the basal ganglia output population **a** over which a target function **f** could be specified. Given the broadly overlapping targets of neighbouring SNr neurons (Deniau and Chevalier, 1992) there could be just one population whose goal is to create a specific trajectory of activity in its targets (Wärnberg and Kumar, 2023).

A second issue is of how control of the dynamic weights can be learnt. The theory outlined here assumes the connections made by the basal ganglia output nuclei have fixed strengths (Figure 2), defining the basis functions, and it is the activity of the output neurons that set the weights of those functions. Unlike traditional views of basis functions with weights that are learnt, these dynamic weights can be adjusted on the fly by input from upstream nuclei of the basal ganglia. Learning when and how to adjust them becomes a question of how feedback from the environment can change the input to the output nuclei. An obvious answer is that environmental feedback is the prediction error signal conveyed by phasic dopamine release in the striatum (Schultz et al., 1997; Bayer and Glimcher, 2005; Rutledge et al., 2010; Hart et al., 2014; Howe and Dombeck, 2016); and that this modulates plasticity at cortico-striatal synapses (Reynolds et al., 2001; Shen et al., 2008; Gurney et al., 2015) to change the input to the basal ganglia’s output nuclei. Extending the theory here to include explicit striatal input and cortico-striatal plasticity would be an interesting line of work.

That striatally-targeted dopamine acts like a reward prediction error and modulates cortico-striatal plasticity are key reasons why the basal ganglia are often viewed as a central part of the brain’s distributed system for reinforcement learning (Ito and Doya, 2011). In this view, the value of a state or action is represented in the cortex, cortico-striatal connection weights, or striatal activity (Houk et al., 1995; Joel et al., 2002; Khamassi et al., 2005; Samejima et al., 2005; Bogacz and Larsen, 2011; Collins and Frank, 2014; Blackwell and Doya, 2023) [see also Elber-Dorozko and Loewenstein (2018) for dissenting evidence], and the downstream basal ganglia circuit enact policy by selecting (Frank, 2005; Humphries et al., 2012; Collins and Frank, 2014) or specifying (Yttri and Dudman, 2016) an action based on that value. As reinforcement learning algorithms place no constraints on how policy is implemented, this view is silent on *how* the basal ganglia circuit selects or specifies action, and silent on the existence of the bottleneck and of any solutions to it. Some forms of reinforcement learning use basis functions to approximate a continuous space of states or actions (Sutton and Barto, 2018). This use of basis functions is distinct from the ideas offered in this paper: here, the basis functions are a property of anatomy, not abstract quantities, and are not explicitly representing anything. Indeed, all theories mapping elements of reinforcement learning algorithms to the basal ganglia could turn out to be false, but if they were false it would not affect the ideas in this paper. The converse may not hold: an open question is whether the re-expansion of basal ganglia output places any limits on mappings of reinforcement learning algorithms to the basal ganglia circuit. Finally, while the dynamic weights idea provides a solution to the problem of the computational bottleneck, it does not explain why this bottleneck exists. Some ideas seem worth pursuing here. Producing spikes makes up much of a neuron’s energy demand (Attwell and Laughlin, 2001; Laughlin, 2001; Sengupta et al., 2010), and so if the high activity rates of the output nuclei are necessary to their function, this could place energetic constraints on their size. Such bottlenecks can also occur as networks evolve when the mapping between their input and output has fewer dimensions than either of the input or the output itself (Friedlander et al., 2015). But, however it arose, the computational bottleneck of basal ganglia output provides a strong challenge to any theories for functions of the basal ganglia.

## Acknowledgements

Though there is a single author’s name on the byline, these ideas represent the synthesis of work from many labs in the vibrant basal ganglia field. They particularly benefited from feedback on my initial presentation of these ideas at the 2016 GRC Basal Ganglia conference, for which I thank Nicole Calakos & Mark Bevan for the invitation to speak, and from the attendees of the 2023 IBAGS meeting. I thank Tom Gilbertson and Bhadra Santhi Kumar for comments on drafts.

This work was supported by the Medical Research Council [grant numbers MR/J008648/1, MR/P005659/1 and MR/S025944/1] and Innovate UK [grant number 10036282].

## Materials and methods

### Code availability

All MATLAB code is available under a MIT License from https://github.com/mdhumphries/Basal_Ganglia_Bottleneck_simulations.

### Estimating the expansion of basal ganglia output

Targets of the mouse SNr were accessed from the Allen Mouse Brain Atlas (Oh et al., 2014) at https://connectivity.brain-map.org/. The Atlas contains seven experiments that had the injection of a fluorescent anterograde tracer targeted to the SNr of the right hemisphere. One experiment (ID:3035444) filled a volume (0.04mm^3^) an order of magnitude smaller than the others, occupying less than 3% of the SNr, and so was not used here. Table 1 gives data on the six retained experiments.

**Table 1:** Allen Mouse Brain Atlas experiments. The proportion of each injection that fell in the SNr was manually scraped from each experiment’s webpage at https://connectivity.brain-map.org/. Proportion of SNr injected used the SNr volume of 0.69 mm^3^ from the Blue Brain Cell Atlas.

| ID | Volume injected ($\text{mm}^3$) | Proportion in SNr | Proportion of SNr injected |
| --- | --- | --- | --- |
| 478096249 | 0.49 | 0.41 | 0.30 |
| 158914182 | 0.54 | 0.29 | 0.23 |
| 478097069 | 0.43 | 0.32 | 0.21 |
| 100141993 | 0.23 | 0.95 | 0.32 |
| 299895444 | 0.36 | 0.94 | 0.51 |
| 175263063 | 0.17 | 0.96 | 0.24 |

Each experiment contained a complete list of all regions in the Allen Common Coordinate Framework V3.0 that had tracer found in them, in both hemispheres, and the volume and density of fluorescent pixels in each region. The Allen Common Coordinate Framework follows a hierarchy from the whole brain, through the grey matter, down to fine-grained subdivisions of nuclei: identified target regions thus overlapped with, or were subdivisions of, others. To find a single, non-overlapping set of targets, I used the list of 295 non-overlapping grey matter regions from Oh et al. (2014), their Supplementary Table 1.

A complete set of neuron numbers for each of those regions in the mouse brain was obtained from Blue Brain Cell Atlas dataset published in Rodarie et al. (2022), their S2 Excel file.

With these data to hand, for each experiment I did the following:

1. Found all targets within the list of 295 unique regions, in both hemispheres
2. Rejected any target region that had a fluorescent pixel density less than some threshold *θ*
3. Looked up the neuron count *nt* in each target region *t* from the Blue Brain Cell Atlas
4. Computed the total potential SNr connections: summed neurons across all *N* retained target regions 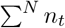
5. Computed an estimate of SNr connections by estimating axonal arborisation in each target: weighted the neuron count in each region by its density *dt* ∈ [0, 1) of fluorescent tracer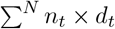.

Thresholding by the density of fluorescence within each target region (step 2) eliminated image noise and other artefacts: without a threshold, all 295 regions contained fluorescent pixels implausibly suggesting the SNr projected to the entire brain (Figure 6). Linear models were fit to the scaling of the total number of target neurons (step 4) and of the estimate of SNr connections (step 5) with the volume of injection in each experiment (fitlm, MATLAB2023a). Extrapolating these models to the volume of the SNr (predict, MATLAB2023a) gave upper and lower bounds on the total number of SNr target neurons. From these were computed upper and lower bounds on the ratio of target to SNr neurons. Computing the upper and lower ratio bounds for a range of density threshold *θ* between 0 and 0.05 in steps of 0.005 revealed they reached an asymptote with increasing *θ* (Figure 1c,d) implying a stable estimate. The asymptote itself was detected as three consecutive bound estimates that changed by less than 1, and was reported as the final of these estimates.

**Figure 6:**
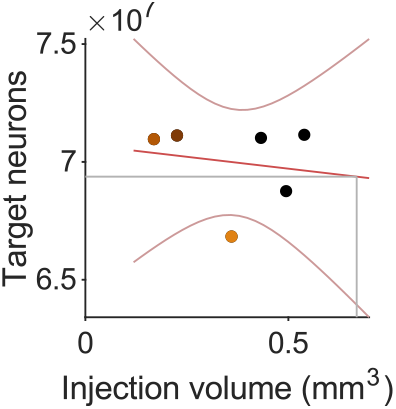
The need for a threshold for noise in tracing experiments. Here I plot the total number of target neurons when including every target region of the SNr with detectable fluorescent pixels in the Allen Mouse Brain Connectivity Atlas experiments. As all or almost all regions had detectable pixels, the total number of neurons is at or close to the total number of neurons in the mouse brain. Also note how the number of neurons does not scale with the injected volume.

As Table 1 shows, three of the experiments had injections that considerably exceeded the bounds of the SNr. Robustness to this limitation was tested in two ways. In the first, the linear model was fit weighting the data points by the proportion of SNr filled in each experiment: this gave the same results for the asymptotic ratios as the unweighted linear model.

In the second, I separately considered each of the three experiments with more than 90% of their injection within the SNr. For each experiment, I first computed the estimated total number of target neurons and the estimate of SNr connections (steps 4 and 5 above). I then computed the estimated number of SNr neurons *Ŝ*within the injection volume, given by

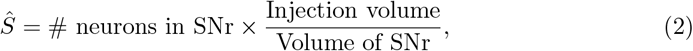

and finally computed the ratio of target neurons to *Ŝ*. The resulting upper and lower bound ratios of target to SNr neuron numbers also reached an asymptote for each of three experiments (Figure 1e,f). The asymptote was computed as above separately for each experiment; the mean of these is reported.

### Using basis functions for expansion

My proposal here stems from the use of basis functions to approximate a target function, so I give a brief account of that here to make the analogy clear.

The target function is *f* (*x*), which can be of any dimension (for example, I use one-dimensional functions in Figures 2 and 4, and two-dimensional in Fig 5). In the most general form of approximating *f* (*x*), we choose some basis function *g*(*x*, Ω), which is defined on some interval *x* and takes parameters Ω, and we obtain the approximation *f* ^′^(*x*) of *f* (*x*) by

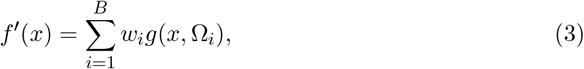

the sum of *B* such basis functions each parameterised by Ω*i*.

Approximating a function typically uses the radial basis functions *g*(*x* − *c, σ*), which are symmetric about their centre *c* and have a single parameter *σ* that defines their width. Gaussians are one type of this class of functions, as shown in Figure 2a. The approximation of *f* (*x*) is then

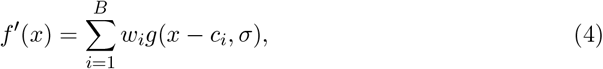

the sum over a set of *B* such basis functions, placed at centres *c*1, *c*2, …, *cB*, and each with its own weight *wi*. Approximating *f* (*x*) then depends on choosing *g*(*x*), setting an appropriate number *B* of these, and finding appropriate weights to minimise some error function *e*(*f* (*x*) − *f* ^′^(*x*)). When using basis functions to fit curves (in one dimension) or surfaces (in two or more dimensions) to sampled data, typically the number and location of sampled data-points respectively determines *B* and the set of *ci*.

Expansion occurs because we want the target function *f* (*x*) to be defined over *n* points in one dimension (or *n* × *m* points in two dimensions, and so on), and typically *B* ≪ *n*. This is especially true when using basis functions to approximate functions across a full range of *n* points using just *B* sampled data points.

### The basal ganglia output as a set of dynamic weights

The theory outlined here is that *g*(*x*) is realised by the strength distribution of an output neuron and the weights *wi* are defined by the activity of that neuron (or equivalently, the strength distribution of a group of output neurons taken collectively, and their collective activity). Then *f* ^′^ is the inhibition function defining the input to every neuron in the target nucleus.

To make this more concrete, consider that there are *b* basal ganglia output neurons that project to target region(s) containing *n* neurons, with *b* ≪ *n*. Then expressing the basis function idea in matrix form, we get:

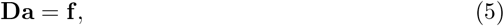

where **a** is the *b*-length vector of basal ganglia output activity, **f** is the *n* length vector of inhibition arriving at the target neurons – the target function in Equation 4 – and **D** is the *n* × *b* matrix that defines the basis functions. For example, for a one-dimensional set of overlapping basis functions that define a ring (cf Figure 4a), each column of **D** is a basis function, shifted circularly across columns.

If the columns of **D** are each a circularly shifted version of the same basis function **v**, and no column can be identical, then the construction of **D** is:

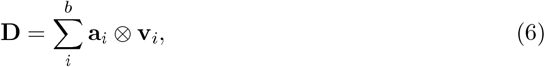

where **v***i* is the *i*th shifted version of **v** and **a***i* is a vector with a 1 at the *i*th entry and 0s at all others. Consequently, Eq 6 defines a full rank matrix, i.e. its columns are linearly independent.

### Recurrent neural network modelling of cortex

The recurrent neural network model of cortex contained *N* = 200 units, 80% excitatory and 20% inhibitory, whose dynamics were given by:

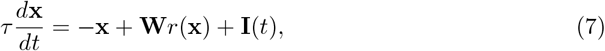

where **x** is the *N* -length vector of unit potentials, *τ* = 20 ms is the network’s time constant, **W** is the *N* × *N* connectivity matrix, *r*() is a function that defines each unit’s firing rate, and **I**(*t*) is an *N* -length vector of inputs to each unit at time *t*. I used here the stabilised supra-linear network (Ahmadian et al., 2013; Rubin et al., 2015; Hennequin et al., 2018) in which the firing rate of the *i*th unit was given by the rectified power-law,

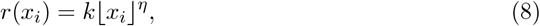

with *k* = 0.1 and *η* = 2. Numerical simulations used forward Euler with a time-step of 0.1ms.

Random connectivity matrices **W** of size *N* × *N* were created by connecting units with probability *p* = 0.1, then randomly assigning columns to be excitatory or inhibitory units. Weights of connections in a column were then defined as 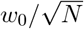 for excitatory connections and 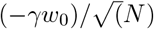 for inhibitory connections, where 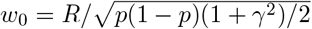, the E:I weight ratio *γ* = 3, and the spectral radius *R* = 10 (Rajan and Abbott, 2006; Hennequin et al., 2014). I tested versions with and without further stability optimisation of the resulting random matrix **W**; further optimisation used the algorithm of (Hennequin et al., 2014) that minimises the maximum real eigenvalue of **W**, resulting in networks whose dynamics recapitulate those of motor cortex (Hennequin et al., 2014; Rodriguez et al., 2024). Similar results were obtained in both random and optimised **W**; results from a random network are plotted in Figure 4. Similar results were also obtained in networks with a 50:50 ratio of excitatory and inhibitory units (not shown).

The vector **I**(*t*) comprised inputs from unmodelled regions of cortex and from thalamus. Background input of **I**(*t*) = 0.1 was applied for *t <* 1000 and *t >* 2000. For time 1000 ≤ *t* ≤ 2000 the inputs stepped up or down from background.

The stepped thalamic input to cortex was assigned to *n* = 0.1*N* randomly chosen entries of **I**(*t*). It was modelled as the vector **T** whose *n* entries were given by **T** = *Tb* − **Da**, where *Tb* = 1 was baseline thalamic activity, **a** was the *b* = 0.25*t* length vector of basal ganglia output activity randomly chosen in [0, 0.4], and **D** was the *n* × *b* matrix of connection weights from basal ganglia output to thalamus. Each column of **D** was thus the basis function defined by each basal ganglia output neuron’s projections to thalamus: here each basis function was symmetric with 1 at the centre and steps down of 0.2 on either side; the basis functions were circularly shifted to uniformly tile the matrix (Figure 4a).

The stepped cortical input was modelled by the remaining 0.9*N* entries in **I**(*t*), which were randomly assigned an initial value in [0, 1]. The stepped thalamic and cortical input were then held constant for 1000 ≤ *t* ≤ 2000.

When simulating the effects of changing basal ganglia output, only the entries of **a** were resampled in each simulation, with all other stepped inputs **I**(*t*) sampled once and then held constant across simulations.

### Superior colliculus grid model

The build-up layer of the superior colliculus was modelled as a *N* × *N* grid of units, with *N* = 20 here. The dynamics of units on the grid were given by

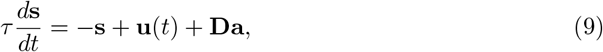

where **s** was the *N* × *N* length vector of collicular unit activity and *τ* = 10 ms was the unit’s time constant. Inputs specifying the saccadic target location were represented in the *N* × *N* -length vector **u**. Input from *b* = 2*N* basal ganglia output neurons was defined by the *b*-length vector of their output **a** and the (*N* × *N*) × *b* matrix **D** that defined the basis functions. The entries of **D** were -1 or 0, set so that each output neuron connected to either an entire row or column of the collicular grid (Figure 5a, bottom).

Colliculus output activity plotted in Figure 5 is given by the rectified linear function *r* = max(0, *s*).

## Appendix: Limits on the number of thalamic neurons reachable by basal ganglia output

The model of basal ganglia output to thalamus adopted in the main text (Figure 2) gives some interesting insights into potential constraints on the divergence of basal ganglia projections to thalamus.

In the model, each of the *b* basal ganglia output neurons has a symmetric distribution of output strengths (main text Figure 2a, inset), which defines its basis function. Given the number of possible discrete strength values *Nb*, the width of this distribution is *NT* = 2*Nb* − 1, which defines the number of thalamic neurons reachable by one basal ganglia output neuron.

Each basal ganglia output strength distribution is centred on a specific thalamic neuron *ti*. Let’s assume the distributions tile the thalamus in a one-dimensional ring to avoid edge effects, and the centres of the distributions are spaced the same number *d* of thalamic neurons apart. The number of possible thalamic neurons in this ring model is then *T* = *d*·*b*.

If we want every thalamic neuron to receive basal ganglia input, then the maximum distance *d* between the centres of the distributions must also be the width of the distributions: *d*max = 2*Nb* − 1. Consequently, the maximum number of thalamic neurons must be *T*max = (2*Nb* − 1)*b*.

But if the basal ganglia output distributions are spaced at distance *d*max then each thalamic neuron receives only one basal ganglia input, and so the basis functions are not overlapping. The distance between basal ganglia output distributions must therefore be *d < d*max. How much smaller would depend on the minimum desired number of overlapping basal ganglia inputs to each thalamic neuron.

Table 2 gives the minimum number of overlapping basal ganglia output neurons in this ring model, for a range of distances *d* between, and resolutions *Nb* of, the basal ganglia output strength distributions. For example, with the centres of the distributions spaced *d* = 3 thalamic neurons apart, and the distributions having *NS* = 6 possible strength values, each thalamic neuron receives input from at least three basal ganglia output neurons.

**Table 2:** Minimum number of basal ganglia inputs to one thalamic neuron. Each entry is the minimum number of inputs each thalamic neuron receives at the specified strength resolution (*Nb*) and distance *d* between strength distributions. These were numerically enumerated, using *b* = 2 · max *Nb*.

| $d$ | $N_b$ | | | | | | | | | | | | | | | |
| --- | --- | --- | --- | --- | --- | --- | --- | --- | --- | --- | --- | --- | --- | --- | --- | --- |
|  | 3 | 4 | 5 | 6 | 7 | 8 | 9 | 10 | 11 | 12 | 13 | 14 | 15 | 16 |  |  |
| 1 | 5 | 7 | 9 | 11 | 13 | 15 | 17 | 19 | 21 | 23 | 25 | 27 | 29 | 31 |  |  |
| 2 | 2 | 3 | 4 | 5 | 6 | 7 | 8 | 9 | 10 | 11 | 12 | 13 | 14 | 15 |  |  |
| 3 | 1 | 2 | 3 | 3 | 4 | 5 | 5 | 6 | 7 | 7 | 8 | 9 | 9 | 10 |  |  |
| 4 | 1 | 1 | 2 | 2 | 3 | 3 | 4 | 4 | 5 | 5 | 6 | 6 | 7 | 7 |  |  |
| 5 | 1 | 1 | 1 | 2 | 2 | 3 | 3 | 3 | 4 | 4 | 5 | 5 | 5 | 6 |  |  |
| 6 | 0 | 1 | 1 | 1 | 2 | 2 | 2 | 3 | 3 | 3 | 4 | 4 | 4 | 5 |  |  |
| 7 | 0 | 1 | 1 | 1 | 1 | 2 | 2 | 2 | 3 | 3 | 3 | 3 | 4 | 4 |  |  |
| 8 | 0 | 0 | 1 | 1 | 1 | 1 | 2 | 2 | 2 | 2 | 3 | 3 | 3 | 3 |  |  |
| 9 | 0 | 0 | 1 | 1 | 1 | 1 | 1 | 2 | 2 | 2 | 2 | 3 | 3 | 3 |  |  |
| 10 | 0 | 0 | 0 | 1 | 1 | 1 | 1 | 1 | 2 | 2 | 2 | 2 | 2 | 3 |  |  |

We see that the minimum number of inputs falls rapidly with increasing distance between distributions. If the distance is greater than 4, then increasing the minimum number of inputs above 2 or 3 requires a large increase in the possible number of strength values. What then should *d* and *Nb* be?

A conservative lower bound on the estimated ratio of thalamic to basal ganglia output neurons is on the order of 5:1 (Figure 1). As the number of thalamic neurons in our ring model here is *T* = *d* · *b*, so this ratio implies *d* = 5. And with that distance between strength distributions, that implies a strength resolution of at least *Nb* ≥ 11 for each thalamic neuron to receive input from at least four basal ganglia output neurons.

